# FastSpeciesTree: Fast and Scalable Species tree Inference

**DOI:** 10.64898/2026.01.20.700630

**Authors:** Jonathan Holmes, Steven Kelly

**Author notes:** ***Corresponding Author*** Name: Steven Kelly, Address: Department of Biology, University of Oxford, South Parks Road, Oxford, OX1 3RB, UK.

## Abstract

The generation of species trees from genomic data is an important step in understanding evolutionary relationships across the tree of life. With the increasing availability of genomic data, species tree inference methods which can easily and rapidly produce accurate species trees from large datasets are required to enable downstream analyses of these resources. Here we present FastSpeciesTree, a fully automated species tree inference method designed to leverage advances in rapid pairwise sequence alignment and scalable phylogenetic inference methods. We show that FastSpeciesTree is able to produce species trees with accuracies equivalent to trees produced through manual-curation and best-practice approaches. We further demonstrate that FastSpeciesTree can produce these high accuracy phylogenies substantially faster than any competitor method. Finally, we show that FastSpeciesTree is capable of building a species tree comprising almost the entirety of the RefSeq databases available proteomes (n = 33285) in less than 12 hours using standard computing resources. FastSpeciesTree is executed as a single command and can be found at https://github.com/OrthoFinder/FastSpeciesTree.

## Introduction

The inference of accurate species trees is of paramount importance to comparative genomics (Majidian, Nevers et al. 2025, Emms, David M., Liu et al. 2025a, Derelle, Philippe et al. 2020). It enables the study and understanding of the phylogenetic relationships between species and provides a platform for studying evolution and diversity across the tree of life (Soltis, Soltis 2003, Meadows, Lindblad-Toh 2017, Genereux, Serres et al. 2020). Given the importance of species tree inference to downstream biological research, multiple methods have been developed for tackling the problem using a variety of different approaches (Zou, Zhang et al. 2024, Dylus, Nevers et al. 2022). These approaches include multi-gene concatenation methods (Gadagkar, Rosenberg et al. 2005), cohort analysis of gene duplications (Emms, David, Kelly 2018) and multispecies coalescence with single or multicopy orthogroups (Zhang, Nielsen et al. 2025a).

Although multiple disparate methods have been generated, each generally relies on the prior construction of orthogroups and the generation of multiple sequence alignments from these orthogroups. The various methods then diverge in the manner in which they utilize these multiple sequence alignments. For example, species tree inference by concatenation utilizes alignments of conserved single copy orthologs which are each assumed to represent the results of evolution along a common ancestral path (Kapli, Yang et al. 2020). However, incomplete lineage sorting can cause the ancestry of individual genes to be incongruent with a single common species tree (Warnow 2015). To overcome this incongruence, tools such as ASTRAL employ a multispecies coalescence model to identify a species tree that maximises the similarity between trees inferred from individual gene multiple sequence alignments (Mirarab, Reaz et al. 2014, Zhang, Scornavacca et al. 2020). Whilst the use of multi-species coalescence models has been shown to accurately infer species trees, the method is ultimately dependent on the requirement for identification of single or multi-copy orthogroups and construction of multiple sequence alignments prior to running an analysis. Thus, the scalability and accuracy of these methods are limited by the underlying scalability of orthogroup inference methods (Emms, David M., Liu et al. 2025b, Emms, David M., Kelly 2019) and in the selection of orthogroups (Swenson, El-Mabrouk 2012).

The availability of low-cost sequence data has led to an increase in the availability of genome data for many species (Sayers, Cavanaugh et al. 2024, Howley, Haas et al. 2025). This increase in data availability has created a new challenge for species tree inference. Specifically, how to infer accurate species trees efficiently from large sequence datasets. With ever improving access to genomic and proteomic data there is a growing need for individuals and research groups to collect and analysis large genomic resources. As a consequence, scalability is becoming increasingly important as computing resources required to infer species trees for these large datasets is not possible for many (Truszkowski, Perrigo et al. 2023, Commichaux, Luan et al. 2024, Pisani, Rossi et al. 2022). Thus, there is a requirement for a species tree inference method which focuses primarily on scalability while providing a high degree of accuracy (Morel, Schade et al. 2022, Zhang, Nielsen et al. 2025b).

Here we present FastSpeciesTree, a species tree inference method designed to infer accurate species trees for tens of thousands of species on conventional computing resources. FastSpeciesTree requires only raw proteome files as input and leverages scalable indexed alignments and phylogenetic inference approaches to rapidly produce highly accurate species trees. We also show that FastSpeciesTree is capable of accurately recreating reference phylogenies across the tree of life. Finally, we demonstrate that FastSpeciesTree is able to outperform a range of alternative species tree inference methods on a broad range of disparate datasets.

## Results

### Implementation and workflow

FastSpeciesTree was designed to infer a species tree from a set of input proteomes files, one per species, with no prior knowledge of species, diversity, or requirement for user processing. FastSpeciesTree is written entirely in python (v3.10.18 <) and is executed as a single command, allowing for simple installation and deployment. The workflow for the method is shown in Figure 1 and described in detail in the methods section. In brief, species tree inference by FastSpeciesTree proceeds in four steps: database preparation and sequence searching (Figure 1A), informative BUSCO (Manni, Berkeley et al. 2021) gene identification (Figure 1B), indexed alignment extraction followed by concatenation and trimming (Figure 1C), and finally phylogenetic tree inference using either the fast or sensitive workflow (Figure 1D). The fast workflow employs VeryFastTree (Piñeiro, Abuín et al. 2020), a parallelised version of FastTree (Price, Dehal et al. 2009) for rapid tree inference, while the sensitive workflow uses IQ-TREE (Minh, Schmidt et al. 2020) on a partitioned alignment file. Each step in the method is fully automated and optimised for scalability and accuracy providing users with the ability to easily infer species trees with minimal input.

**Figure 1.**
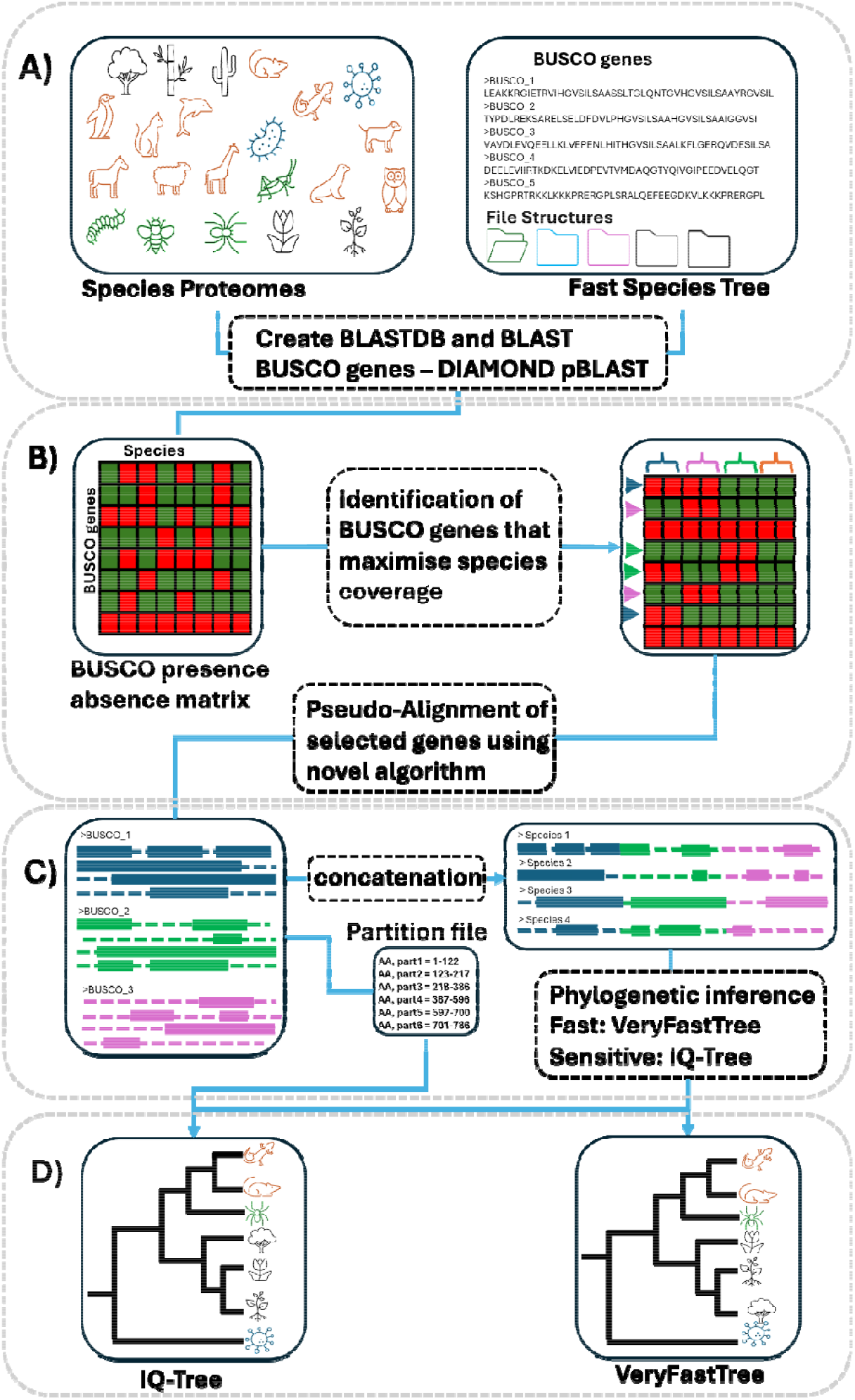
The FastSpeciesTree workflow. FastSpeciesTree consists of 4 steps – A) database preparation and sequence searching, (B) informative BUSCO gene identification, (C) indexed alignment extraction and concatenation, and (D) phylogenetic tree inference.

### Benchmark datasets and comparator methods

In order to assess the accuracy of FastSpeciesTree we compared its output to a range of reference species trees inferred from real data. The reference species trees were selected from a set of highly cited research papers (Zhu, Mai et al. 2019, Liu, Steenwyk et al. 2024, Shen, X., Opulente et al. 2018), which were taken to provide highly-accurate and expert-curated phylogenies that could serve as suitable benchmarks. Wherever possible, the original proteome files were used, however, where proteome files were not supplied in the original study the equivalent proteomes were downloaded from NCBI.

For each benchmark dataset, FastSpeciesTree was run on the input proteomes using its two different workflows: fast (VeryFastTree) and sensitive (IQ-Tree). To provide a comparison of the accuracy and performance characteristics of FastSpeciesTree to competitor methods, three direct comparator methods were also run on the same datasets. These methods were selected for their diverse methodological approaches to species tree inference and comprised Feature Frequency Profile (FFP) (Sims, Jun et al. 2009), busco2phylo-nf (https://github.com/lstevens17/busco2phylo-nf) and Phyling (Tsai, Stajich 2025). FFP is an alignment free method that infers a phylogenetic tree from a supermatrix of l-mer abundances. Busco2phylo (Warnow 2015, Mayeur, Leyhr et al. 2024, McHale, Mulhair et al. 2025a, Mulhair, Crowley et al. 2023, McHale, Mulhair et al. 2025b) uses hidden Markov models computed from alignments of BUSCO genes to identify single copy orthologs within the input proteomes which are subsequently extracted and subject to multiple sequence alignment, gene tree inference, and species tree inference with ASTRAL-Pro3. Phyling, also uses hidden Markov models computed from alignments of BUSCO genes to identify conserved genes which are then aligned and subject to tree inference. In all cases, the published phylogeny was assumed to be true, and the accuracy of all methods was evaluated by comparison to these phylogenies.

### FastSpeciesTree achieves a high degree of topological accuracy on real world datasets

The *opisthokonts* are a group of eukaryotes which contain metazoa and fungi (del Campo, Mallo et al. 2015, Torruella, de Mendoza et al. 2015). Previous expert phylogenetic interrogation of this group inferred a species tree for 347 species using three different approaches (Liu, Steenwyk et al. 2024). In brief, the three approaches involved: the curation of single copy orthologs using either the BUSCO database of single copy orthologs, the single copy orthologs generated using OrthoFinder (Emms, David M., Liu et al. 2025b), or the single copy orthologs curated from Tikhonenkov (Tikhonenkov, Mikhailov et al. 2020). Following ortholog inference, the multiple sequence alignments were manually curated and concatenated and used to infer a species tree using IQ-TREE2. This workflow represents a common approach that is used in many studies to generate species trees (Segata, Börnigen et al. 2013, Li, Steenwyk et al. 2021, Feng, Wan et al. 2025, Hug, Baker et al. 2016).

To determine how FastSpeciesTree and the other comparator methods in each method was run on the set of 347 proteome files above using 32 cores. FastSpeciesTree took 14 minutes to infer the species tree in Fast mode and 14 hours and 23 minutes in sensitive mode. In contrast, FFP took 16 hours, Phyling using concatenation took 28 hours with IQ-Tree (sensitive) and 32 hours with FastTree (fast), and busco2phylo-nf took 194 hours to complete.

Two metrics were used to quantify the topological similarity between the inferred species trees and the reference species trees. These comprised the generalized Robinson–Foulds (gRF) distance and the quartet distance. The gRF distance assesses topological similarity based on individual bipartitions, and the quartet distance evaluates the proportion of identical four-taxon subtrees. Across both metrics, FastSpeciesTree performed comparably to the other evaluated methods. The sensitive workflow of FastSpeciesTree achieved a mean gRF of 0.07 and a mean quartet distance of 0.02. There was no statistically significant difference in topological distance between FastSpeciesTree (sensitive), busco2phylo-nf, and Phyling (sensitive) for either gRF (Figure 2A: group A) or quartet distance (Figure 2B: group A).

**Figure 2.**
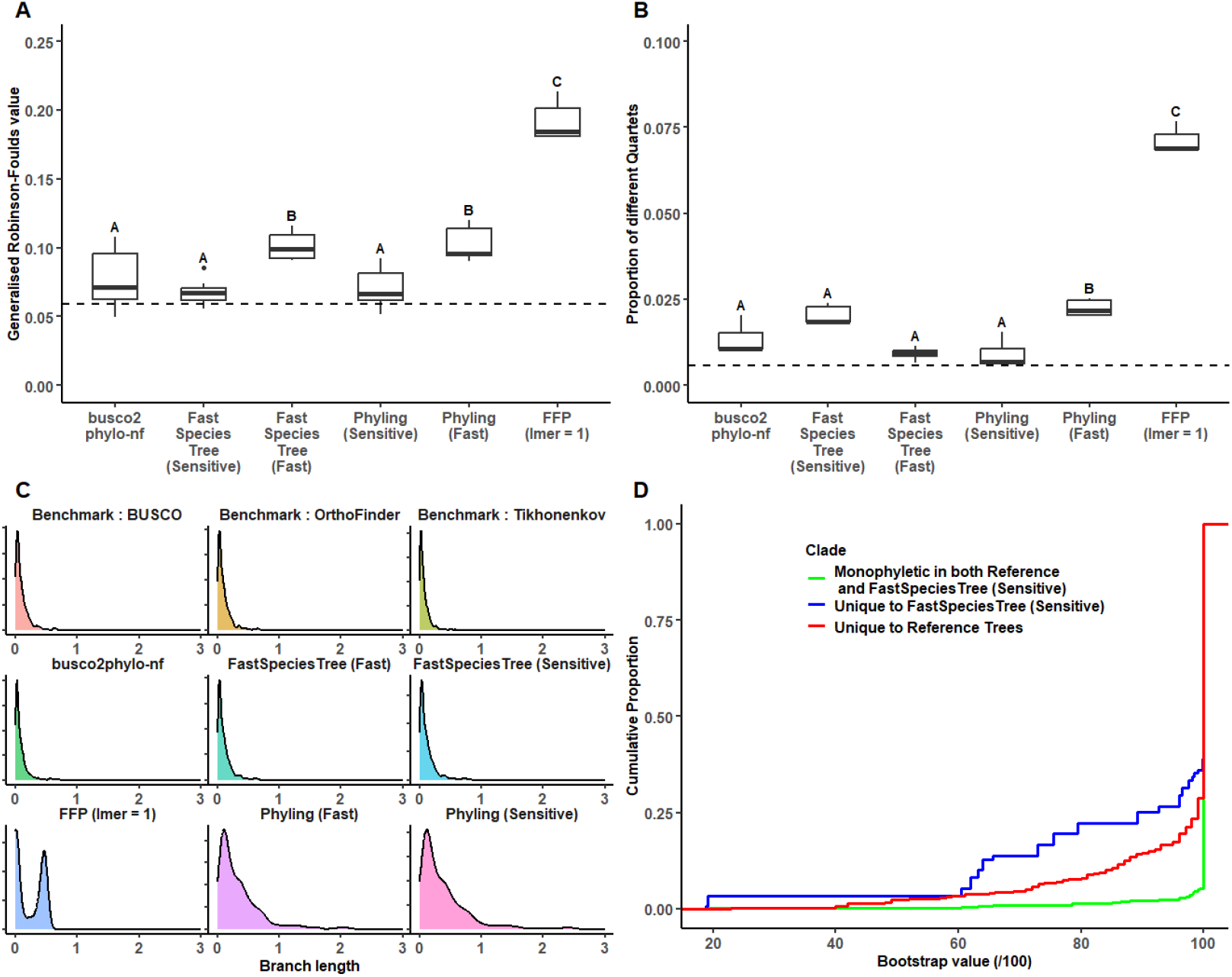
Comparison of the accuracy of the species tree inference methods on *Opsithokont* dataset. A) The generalised Robinson-Foulds (Llabrés, Rosselló et al. 2021) (gRF) distance score the inferred and reference trees. B) The quartet distance between the inferred and reference trees. C) The branch length distributions of the inferred and reference trees. D) The discrepancy between topology as a function of bootstrap support value. Letters above each boxplot indicate statistically significant differences among groups (one-way ANOVA followed by Tukey HSD post-hoc test, *p* < 0.01). In plots A and B the dashed line indicates the level of internal discordance between the reference trees.

The fast workflow of FastSpeciesTree inferred a species tree with a gRF score comparable to that obtained by Phyling using FastTree (Figure 2A: group B and Figure 2B: group B). However, its quartet distance was approximately twofold lower than that of Phyling (fast), indicating improved recovery of the topology of the tree. In contrast, the other alignment-free approach, FFP, exhibited the highest scores across both gRF and quartet distance metrics, significantly higher than all other tools tested. Thus, FastSpeciesTree produces trees with equivalent or higher topological accuracy then comparator methods.

### Topological discordance is primarily localised within poorly supported partitions

To assess whether the differences that were observed between FastSpeciesTree and the reference trees corresponded to partitions with low support values, the support values for partitions that were common or unique to the test and reference trees were analysed (Figure 2D). Equivalent analyses were conducted for FastSpeciesTree (fast) and busco2phylo-nf (Supplementary Figure 3). Clades that were common to the reference trees and FastSpeciesTree exhibited high support values, with the majority achieving 100% support by both methods. In contrast, partitions that were unique to either the reference tree or FastSpeciesTree showed a diverse distribution of support values (Figure 2D). This indicates that regions of topological discordance between FastSpeciesTree and the reference tree are associated with weakly supported partitions in both trees. This discordance with weakly supported partitions is the cause of the differences measured by the gRF and quartet distance scores (Figures 2A and 2B). Similar trends were observed for busco2phylo-nf and FastSpeciesTree (fast) (Supplementary Figure 3). Thus, topological discordance between FastSpeciesTree and the reference tree is primarily contained within poorly supported partitions.

### FastSpeciesTree produces species trees with accurate branch lengths

In addition to topology, the lengths of the branches in a phylogenetic tree are an important property of the tree. For example, branch lengths in species trees are commonly used to estimate divergence times (Tiley, Flouri et al. 2023, Schwartz, Mueller 2010). To assess whether the species tree inference methods produced accurate branch lengths the branch length distributions from each method were compared against those obtained from the three reference tree methods (Figure 2C). FastSpeciesTree, using both workflows, exhibited mean branch length distributions that closely matched those of the reference trees (Figure 2C). Similarly, busco2phylo-nf also produced branch length distributions that were similar to those observed in the reference trees. In contrast, Phyling (fast and sensitive) produced substantially longer branches with an approximately threefold overestimation of evolutionary distance between species (Figure 2C). FFP displayed a bimodal branch length distribution, with one subset comparable to the reference trees and a second peak approximately fivefold greater than the reference mean (Figure 2C). Thus, in addition to high topological accuracy, FastSpeciesTree produces trees with accurate branch lengths.

### FastSpeciesTree also performs well over large evolutionary distances

While FastSpeciesTree performed well on the Opisthokont test above, the crown divergence of this group is only ∼1.3 billion years ago (Auxier, Dee et al. 2019) and thus represents a subsample of the evolutionary distances to which FastSpeciesTree could be applied. To provide a test that encompassed a broader evolutionary divergence, a benchmark dataset comprising a large set of species that encompass the root of the tree of life was evaluated (Zhu, Mai et al. 2019). This reference phylogeny consisted of 9906 bacterial and 669 archaeal species. In brief, Zhu et al (2019), generated a reference phylogeny for these species using two different approaches. First, utilising multicopy gene families and ASTRAL-MP, second using single copy genes and concatenation of multiple sequence alignments. Species trees for this dataset were inferred using FastSpeciesTree and the comparator methods above using 32 CPU cores. Phyling was run using the BUSCO prokaryote dataset to identify genes. The IQ-Tree based methods of FastSpeciesTree, Phyling and busco2phylo-nf, were unable to complete species tree inference using 32 CPUs in a reasonable timeframe or failed to start. However, Phyling in its consensus mode using ASTRAL-Pro3 with FastTree completed the analysis in 110 hours. Phyling in its concatenation mode it required 249 hours to complete. FFP also successfully inferred a phylogeny, requiring 210 hours of computation. In contrast, FastSpeciesTree using the fast workflow inferred the species phylogeny in one hour and 19 minutes. Thus, FastSpeciesTree was able to compute the large species tree ∼84 times faster than the closest comparator method.

The topological similarities between the inferred trees and the reference phylogenies of Zhu *et al* (2019) were quantified using the generalized Robinson–Foulds distance as above. As the available algorithms for computing gRF metrics are unable to handle large phylogenetic trees, a bootstrap consensus gRF score was evaluated by sampling random subsets of 1,000 taxa from both the test and reference trees and evaluating the gRF scores obtained from comparison of these sub-sampled trees. This subsampling procedure was repeated 1,000 times, to provide an ∼100-fold coverage for each species in the dataset. This analysis revealed that both Phyling and FastSpeciesTree produced species trees with similar gRF values. Phyling in its concatenated method was able to produce the species tree that most closely matched the reference phylogeny created by Zhu et al (2019), for both the concatenated reference tree (Figure 3A) and the consensus tree (Figure 3B). Both methods produced species trees that were substantially more accurate than FPP (Figure 3A and 3B). Zhu et al (2019) utilised PhyloPhlAn as a database for its gene selection, a database created for microbial taxonomic placement. In a similar way Phyling is only able to identify genes found in a single BUSCO schema, in this case the prokaryotic schema. Meaning that the species tree generated by both Zhu et al (2019) and Phyling were both generated using predominantly bacterial genes, rather than a combination of genes from both bacterial and archaea databases as with FastSpeciesTree.

**Figure 3.**
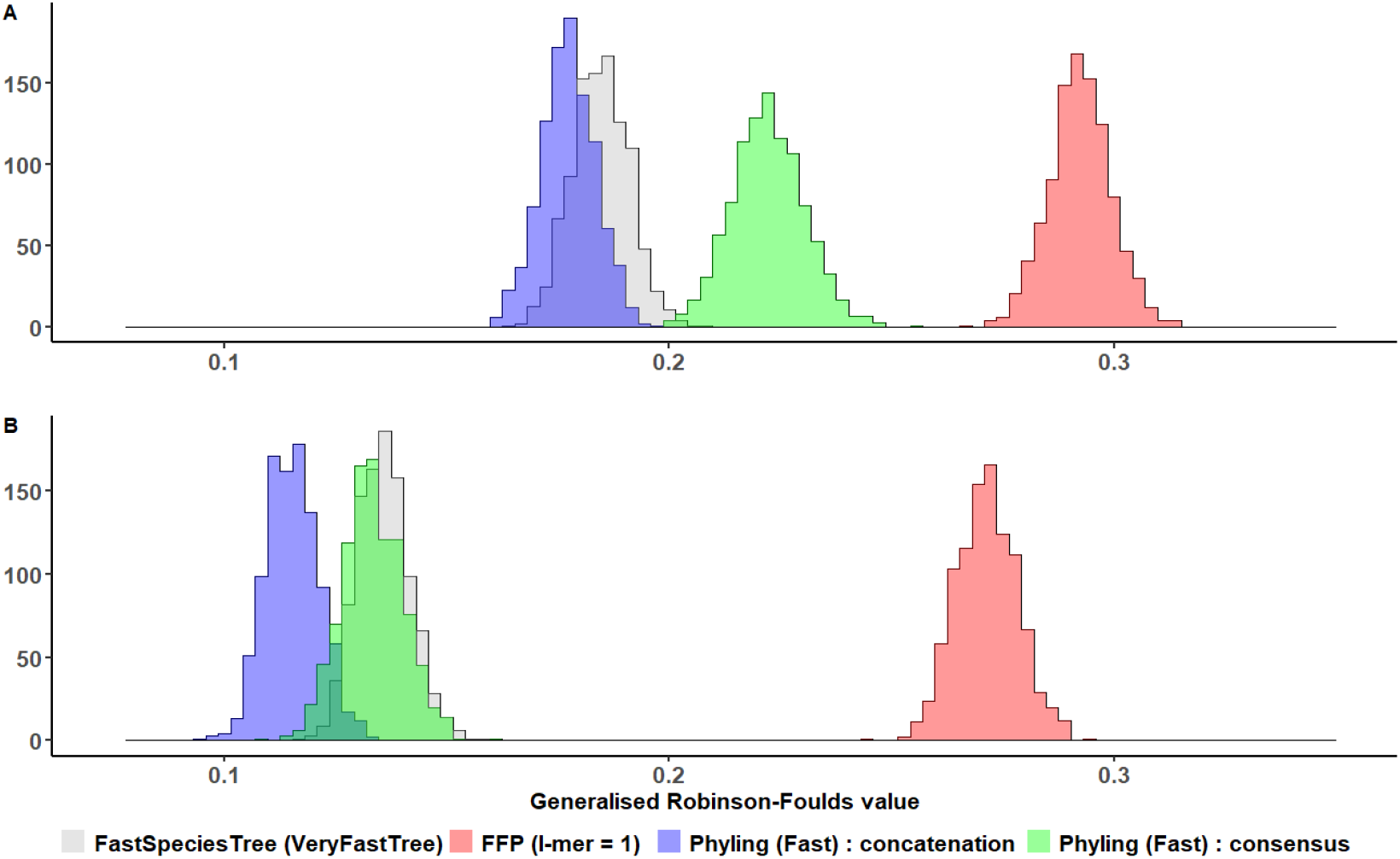
Topological accuracy of the species tree methods on a large bacterial and archaeal dataset. A) Comparison of the different methods to the concatenated alignment-based reference phylogeny. B) Comparison of the different methods to the consensus tree-based reference phylogeny.

### Benchmarking the scalability of FastSpeciesTree

Given that FastSpeciesTree was able to infer accurate species trees over large a range of phylogenetic distances, we next sought to determine how FastSpeciesTree scales with increasing number of species. To do this, a reference dataset of 330 species of budding yeast (Shen, X., Opulente et al. 2018) was subject to species tree inference with subsets of species sampled in increasing powers of 2 (n = [4, 8, 16, 32, 64, 128, 256]). Each subsampled dataset was then subject to phylogenetic tree inference using the cohort of methods using 32 CPU cores, and the runtime characteristics (Figure 4A) and distribution of topological distances relative to a reference phylogeny (Figure 4B) were recorded. Across all dataset sizes tested, FastSpeciesTree in its fast implementation was the fastest (Figure 4A), with an inference time between 20-500-fold faster than the most similar comparator methods (Phyling [fast] and FFP) depending on input dataset size. Similarly, FastSpeciesTree in its sensitive implementation was between 1.5 and eight times faster than comparable sensitive methods (Phyling: sensitive and busco2phylo-nf) depending on input dataset size (Figure 4A). Notably, in the case of Phyling, the runtime advantage of FastSpeciesTree increased with increasing sample size.

**Figure 4.**
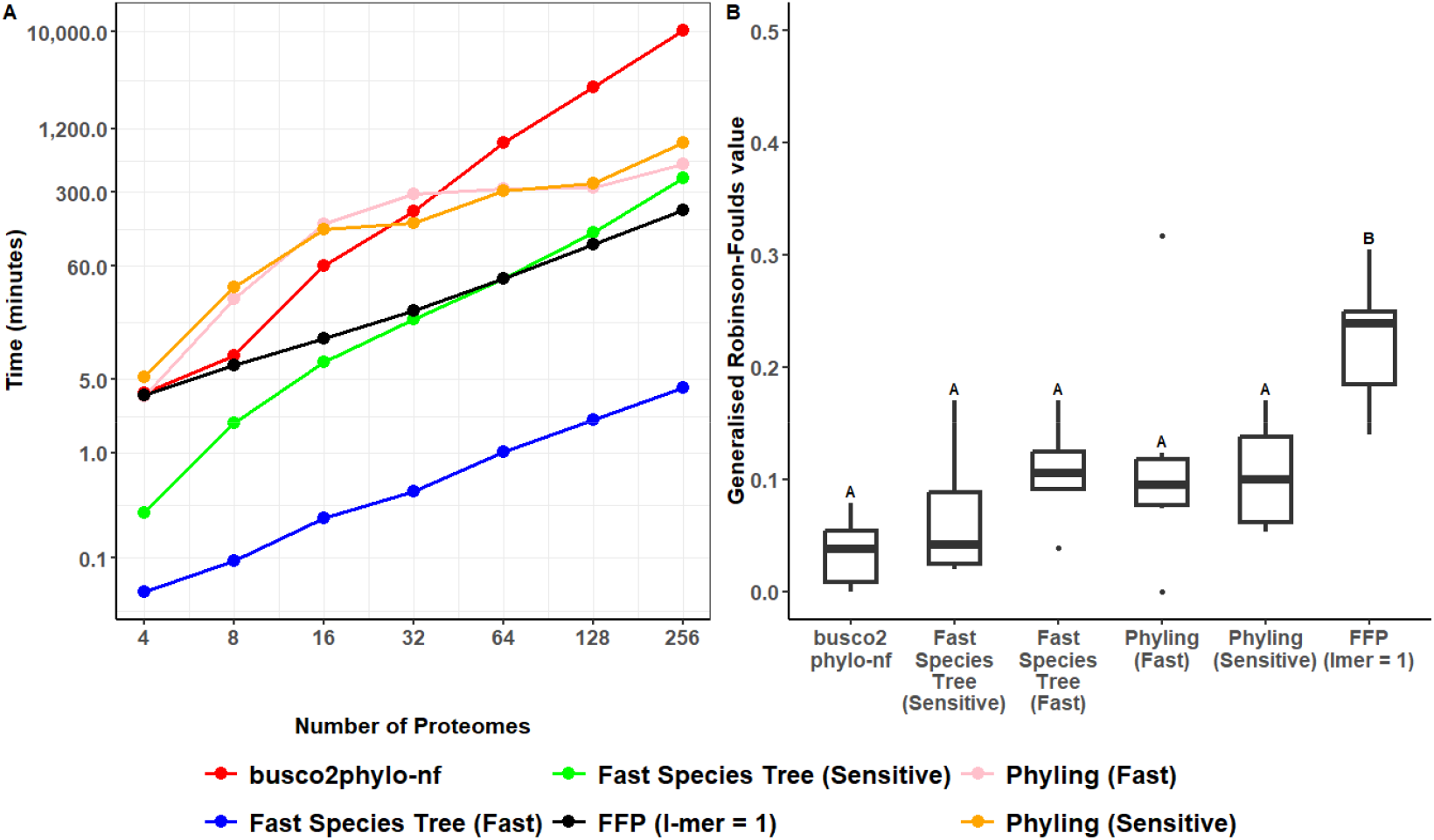
Performance benchmarking for runtime (A) and accuracy (B) on species tree inference methods with increasing dataset sizes on a budding yeast reference phylogeny. All methods were run with 32 cores.

FastSpeciesTree in its sensitive implementation also demonstrated strong performance, consistently outperforming similar comparator pipelines. Specifically, it was between two-fold and 25-fold faster than Phyling with IQ-Tree and busco2phylo-nf across the range of dataset sizes tested (Figure 4A). While the absolute runtime differences were smaller than those observed for the fast workflow, FastSpeciesTree was able to achieve the 256 taxa phylogenetic species tree 2-fold faster than Phyling (fast) and 25-fold faster than busco2phylo-nf without any compromise to tree inference accuracy.

Among the comparator methods, FFP emerged as the second fastest overall, running approximately 20–30 times slower than FastSpeciesTree in fast mode but faster than all remaining pipelines. However, FFP achieved a significantly higher topological distance (Figure 4B) than all other methods tested, and therefore is substantially less accurate than the other methods. In contrast, FastSpeciesTree (both fast and sensitive modes), Phyling, and busco2phylo-nf all achieved broadly comparable levels of accuracy across dataset sizes. Although busco2phylo-nf yielded the lowest overall topological distance relative to the reference tree, its performance was not significantly different from that of FastSpeciesTree in either mode. Importantly, busco2phylo-nf exhibited poor scaling behaviour, with runtimes increasing steeply as dataset size grew, effectively limiting its applicability to smaller datasets. FastSpeciesTree (senstitive) was able to out compete Phyling using FastTree for both runtime (Figure 4A) and accuracy (Figure 4B) with a lower average gRF achieved across the sample sizes.

The discordance patterns observed in this yeast benchmark (Figure 4B) recapitulate those obtained using the Opisthokont benchmark dataset (Figure 2A). In both cases, FFP performed poorly, yielding trees with significantly higher topological distances relative to reference phylogenies than any other method. Notably, in this yeast dataset, FastSpeciesTree and Phyling using their fast workflows did not produce significantly higher discordance (Supplementary Figure 4) than the sensitive based workflows. Thus, FastSpeciesTree is able to create species trees as accurately as alternative methods on reference datasets while simultaneously being able to outcompete these methods for runtime in both its fast and sensitive workflows.

### Demonstrating the utility of FastSpeciesTree on large datasets

To demonstrate the utility of FastSpeciesTree for large species tree inference tasks, the ensembl (Hubbard, Barker et al. 2002) [Accessed: 19/05/2025] release was downloaded and subject to species tree inference. This dataset comprised 31,331 bacteria, 367 metazoa, 1,427 fungi, and 216 plants, totalling 33,341 proteomes. FastSpeciesTree was run with 32 cores using the fast workflow. The species tree was inferred in ∼12 hours (41184 s) with a concatenated alignment of 353 genes (Figure 5). No other species tree inference tool, other than FastSpeciesTree, was able to infer a tree for this dataset. The comparator method that got closest to completing was Phyling (fast). Phyling was able to perform its sequence search phase in ∼6 hours but was unable to create a phylogenetic tree using all 129 detected genes from the Eukaryotic BUSCO schema. However, it was possible to produce a phylogenetic tree using a reduced set of 20 genes, selected using Phylings filter workflow, in two days using FastTree (Supplementary Figure 5). Additionally, FFP one of the fastest methods (Figure 4A) was also unable to infer this phylogeny due to requiring ≥512Gb of RAM to create the distance matrix needed to infer the tree.

**Figure 5.**
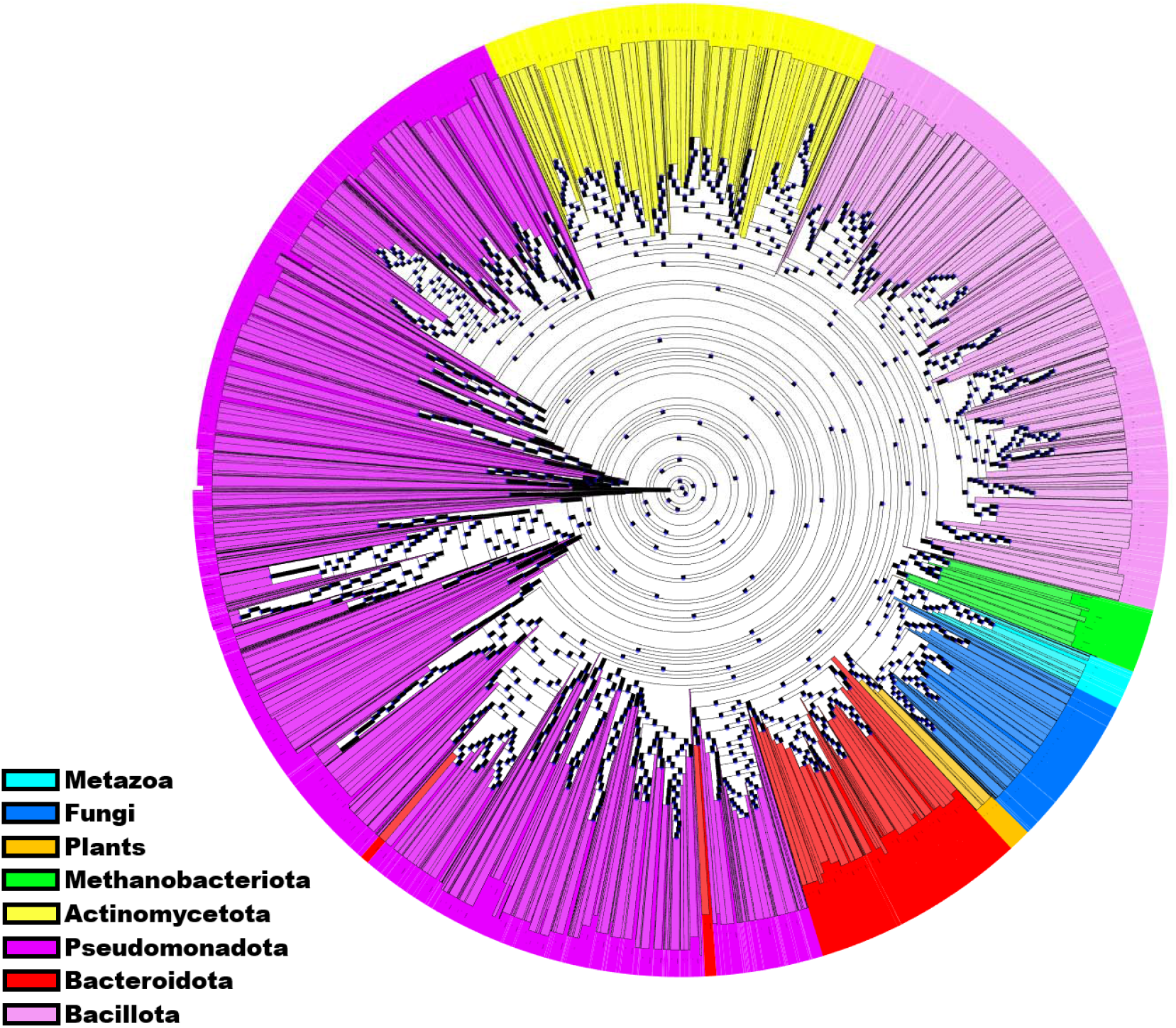
A phylogeny of the complete ensembl release (19/05/2025) generated using FastSpeciesTree (fast). Major bacterial phyla along with Eukaryotes are highlighted. Annotations for the plants, fungi and metazoa are based on the ensembl dataset, while the bacteria which make up the majority of species (94 %) were highlighted based on major phylum, derived from BacDive (Schober, Koblitz et al. 2025) annotations and visualised with ETE4 (Huerta-Cepas, Serra et al. 2016) showing the domains of Fungi, Plants, Metazoa and the major bacterial phyla of *Actinomycetota, Psuedomonadota, Bacillota, Bacteroidota* and *Methanobacyeriota*.

Although no ground truth or widely accepted phylogeny exists for this diverse set of species, several approaches can be used to evaluate the quality of the inferred tree. Notably, we expect major domains of life and key bacterial phyla to form monophyletic clades. Consequently, clades that intermix disparate groups—such as Plants and Metazoa—are considered erroneous. To assess this, we traversed the unrooted trees inferred by FastSpeciesTree and Phyling bidirectionally, evaluating each node for clades for mis-placed taxa (Figure 4). The resulting clades were ranked by size, with the largest clade for each group representing the maximum fraction of that group’s taxa recovered in a single, high-purity clade within the species tree (Supplementary Table 1). Overall, the tree produced by FastSpeciesTree achieved an average of 99.7% consistent taxon placement across the eight major groups. In contrast, Phyling attained only 75% consistent taxon placement. It particularly underperformed on Metazoa, placing just 31% of the 367 taxa (i.e., 116 taxa) in a monophyletic group, whereas FastSpeciesTree placed 100% (Figure 5, light blue metadata). Thus, FastSpeciesTree is a viable approach for producing accurate species trees for large scale analyses.

## Discussion

Species tree inference underpins much of contemporary comparative genomic research (Shen, F., Qin et al. 2023, Alföldi, Lindblad-Toh 2013, Zoccarato, Sher et al. 2022). With the increasing availability of genome data, it is essential that methods are developed that can accurately and efficiently analyze these resources to maximise their value. Here we present FastSpeciesTree, a method that makes use of widely conserved genes, fast pairwise alignment tools, and scalable phylogenetic inference methods to provide an accurate and rapid method for inferring species trees.

FastSpeciesTree was designed with usability and speed as a priority. The packaging of the workflow into a single Python script allows for flexibility and simple downloading along with minimal package requirements for installation. This provides accessibility for users with limited experience and ease of integration into existing comparative genomics workflows. FastSpeciesTree gains its scalability primarily through the use of pseudo-alignments generated from DIAMOND pairwise sequence similarity searches and the optimisation of much of the workflow to run using parallel cores. This optimisation allows FastSpeciesTree to infer phylogenies for large datasets in-user friendly timeframes without complex data pre or post processing requirements.

Using a variety of different test cases (*Opisthokonts, Fungi*, Bacteria and Archaea, and a large “tree of life” dataset comprising every species available in ensemble) we demonstrate that FastSpeciesTree is capable of generating species trees more rapidly than comparator methods with equivalent accuracy to best practice approaches on expert curated datasets. We further demonstrated that where differences occurred between the trees produced by FastSpeciesTree and the reference trees, that these primarily occurred at branches that received poor support in both trees. Collectively, these results indicate that the trees produced using FastSpeciesTree accurate and suitable for a broad range of uses in comparative genomics.

Multiple disparate methods that utilise different approaches to species tree inference were used to provide a comparison with FastSpeciesTree. While some showed similar accuracy, none were as scalable as FastSpeciesTree with methods varying between five and 1,000 times slower than FastSpeciesTree depending on the input dataset. FastSpeciesTree, therefore, provides users with a fast, scalable, and accurate species tree inference method. It is important to note that FastSpeciesTree is not intended as a replacement for expert characterisation of species trees. Instead, FastSpeciesTree is provided as a scalable and easy to use method that can inform this process, and that facilitates easy construction of accurate species trees for a variety of different uses. For example, it is envisioned that the method will facilitate production of species trees for large repositories of sequence data and large-scale comparative genomics projects.

## Methods

### Creation of an optimal BUSCO reference gene set

To generate a reference gene set for use in sequence similarity searches the complete set of BUSCO (Manni, Berkeley et al. 2021) genes that are reportedly present in ≥ 90% of species across the three domains of life were downloaded from BUSCO. A single query sequence representing each BUSCO gene was generated using hmmemit (Finn, Clements et al. 2011) to produce a plurality-rule consensus sequence for each hidden Markov model representation of the BUSCO gene for the eukaryotic, bacterial and archaeal schemas. Hidden Markov models for BUSCO genes that produced significant sequence similarity scores against other BUSCO genes were removed from consideration (a cross-reactive e-value threshold of 0.01), to prevent erroneous mis-assignment of genes between BUSCO groups. This resulted in a set of 574 reference BUSCO gene sequences (henceforth termed reference BUSCO genes) that were subsequently used in all downstream analyses.

### Sequence similarity searches

The complete set of input user proteomes are first subject to DIAMOND (Buchfink, Reuter et al. 2021, Buchfink, Xie et al. 2015) index construction. Following this, the set of 574 reference BUSCO genes constructed above are then queried against these proteomes using DIAMONDs BLASTp and the complete set of High-scoring Segment Pairs (HSP’s or “hits”) better than the e-value threshold of 0.01 are retained for subsequent analysis.

### Informative BUSCO gene identification

To select the set of reference BUSCO genes that produce hits for use in phylogenetic analysis, the hits are converted into a binary matrix describing the presence and absence of a hit for each species in the input dataset. The binary presence matrix is then scaled to normalise the mean and standard deviation of each row, using Sklearn’s StandardScalar (Pedregosa, Varoquaux et al. 2018). K-means clustering is then performed across the matrix with up to 20 possible clusters considered. The number of clusters is estimated using a silhouette score, an average measure of how well each object fits into a cluster, with a minimum threshold required for the proteomes to be clustered (≤ 0.25). The number of K-means clusters with the highest silhouette score above the minimum threshold is then used to subdivide the species of interest into groups that share the same sets of BUSCO genes. This best representative BUSCO schema for each given cluster is determined by the total maximum score of that schema, combining overall presence and score of the DIAMOND hits. The BUSCO genes for each cluster that achieve ≥ 70% presence threshold are then selected for subsequent phylogenetic analysis (Liu, Steenwyk et al. 2024). This clustering step is an important section of the workflow, as a requirement to sample genes with high presence across the entire input dataset can cause the analysis to fail to run. For example, in a dataset containing an equal number of Bacteria, Archaea and eukaryotes there is unlikely to be sufficient genes present in 70 % of all input proteomes to facilitate analysis. While a cluster may only be represented by a single BUSCO schema, all hits across all genes are still recorded and used when that gene is selected for by another cluster, ensuring that alignments across these domains of life are possible.

### Construction of the pseudo-multiple sequence alignment

Once the set of BUSCO genes that achieve the requisite coverage thresholds in the input species dataset are identified (Figure 1B), the observed indexed pairwise alignments are then used to construct a concatenated multiple sequence (Figure 1C). Importantly, this step bypasses the requirement to construct a multiple sequence alignment as it leverages the indexed positional information produced by DIAMOND during the sequence similarity search. Specifically, this is achieved by identifying the highest scoring sequence within the input proteome of a given species and retrieving all additional High-Scoring Segment Pairs between this sequence and the reference BUSCO gene. Following this, an optimal pairwise alignment between the BUSCO refence gene and all High-Scoring Segment Pairs originating from the same input gene is constructed using the DIAMOND indexed High-Scoring Segment Pairs information. Sequence positions in the input gene which are not included in an aforementioned High-Scoring Segment Pair, are not included in this pairwise alignment. Similarly, positions in the reference gene that have no corresponding aligned position in the High-Scoring Segment Pair results are represented as a gap character in the reconstructed optimal pairwise alignment. This reconstruction process produces an amino acid sequence that is exactly the same length as the BUSCO reference gene, where each position is either a gap or the amino acid from the identified best scoring gene for a given species. As a corollary of this reconstruction process, the set of reconstructed sequences from each species comprises a multiple sequence alignment of high confidence aligned positions, and thus bypassing the requirement to independently construct a multiple sequence alignment. The complete set of reconstructed alignments are then concatenated for a single contiguous concatenate multiple sequence alignment. Alignment trimming is then performed on the concatenated alignment only in fast mode, as IQ-Tree ignores these sites, to reduce conversed and non-phylogenetically informative sites. During this process, the partition file which contains information about the gene partitions within the concatenated alignment is updated to reflect the reduced number of alignment columns.

### Species tree inference

Following construction of the multiple sequence alignments above, FastSpeciesTree follows one of two different analysis routes dependent on the user’s preference. Specifically, the concatenated alignment is passed to one of two supported tree inference software (Figure 1D): VeryFastTree (4.0.5) or IQ-TREE (3.0.1). Either tree inference software can be selected providing flexibility for scalability (VeryFastTree) or increased accuracy (IQ-TREE). VeryFastTree, a parallelised version of FastTree, takes the concatenated pseudo-alignment as input producing a tree in Newick format. Alternative, if IQ-TREE is selected, then IQ-TREE is then run on the concatenated sequence file using an additional partition file that is generated during the alignment construction step which describes the partitions within the concatenated sequence alignment. A reduced set of available models are used by default, these are: LG, JTT, Q.BIRD, Q.MAMMAL, Q.INSECT, Q.PLANT, Q.YEAST, additionally only I, G and I+G are considered as model rates. The default IQ-TREE runtime options are set to be 1,000 ultrafast bootstraps and 1,000 replicates for SH approximate likelihood ratio test. All necessary input files are stored, should the user wish to re-run the analysis using alternative IQ-TREE models or options, or any other phylogenetic tree inference software which they may prefer.

### Tree comparison and accuracy evaluation

FFP was run using an l-mer length of one. Phyling was run using a relevant BUSCO schema with all identified genes used to build the consensus sequence, unless stated otherwise, with the species tree built with FastTree (fast) and IQ-TREE (sensitive). Busco2phylo-nf was run using a presence threshold of 70 %, based on available data from the reference phylogenies (Liu, Steenwyk et al. 2024) and the cutoff used in FastSpeciesTree, the NextFlow config was modified to run locally using a maximum of 32 CPUs.

In all cases, topological comparisons were made using the generalised Robinsons Foulds (gRF) distance with TreeDist (Smith 2020) and quartet distance (Smith 2022). These two distinct measures quantify the dissimilarity of clades or quartets between two trees with values of zero, indicating identical trees for both gRF and quartet distance. Where there were discrepancies in the number of taxa between methods evaluated then non-overlapping leaf tips were pruned. For the *opisthokont* tree, the “rogue taxa absent” species trees for each of three models used by IQ-TREE (CAT, LG, C60) was taken as the reference tree.

The impact of low support partitions on tree topology was performed for those methods where a support value for each node is provided, FastSpeciesTree and busco2phylo-nf. Partitions were divided into high and low support clades and compared to the reference tree. Each clade was then searched in the reference tree and flagged as either monophyletic and non-monophyletic and its support value recorded as monophyletic in both the reference and test tree, unique to the test tree or unique in the reference tree with non-monophyletic clades indicates incongruence in the placement of species in the tree.

In addition to topological comparisons to the reference phylogenies with gRF and quartet distance, the species trees were also evaluated by their ability to recapture the correct branch length distributions of the reference trees. To assess this, the distribution of edge lengths were collected for all branches in the tree and compared directly between the comparison method and reference phylogenies.

## Author information

JH and SK developed the method. JH implemented software. JH and SK contributed to the analysis. JH and SK wrote and edited the manuscript.

## Ethics declaration

SK is co-founder of Wild Bioscience Ltd and an employee of Ellison Institute of Technology, Oxford Limited.

## Data Availability and Implementation

FastSpeciesTree is available at https://github.com/OrthoFinder/FastSpeciesTree.

