## Supplemental File 1 for "FastSpeciesTree: Fast and Scalable Species tree Inference"

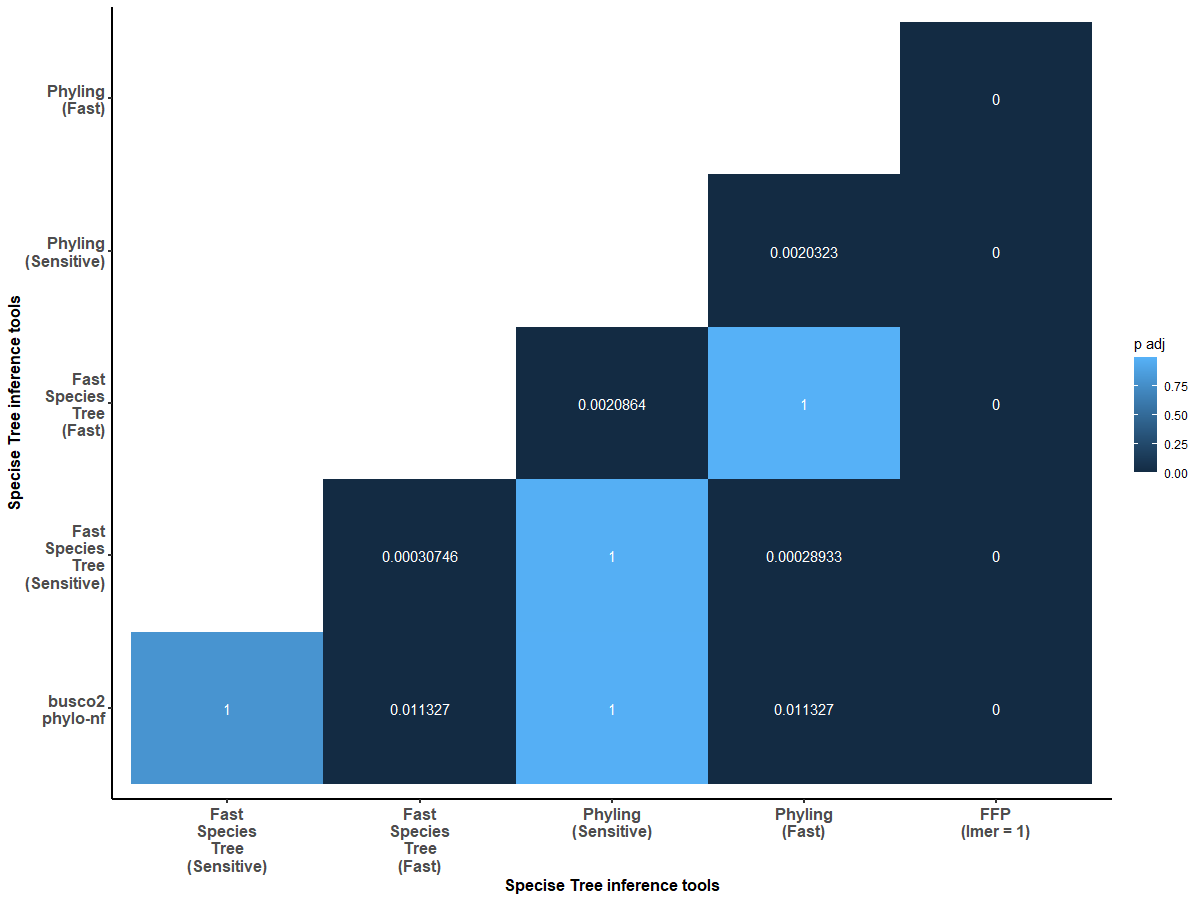

**Figure 1.** The statistical similarity of generalised Robinson-Foulds distance between 4 species tree inference methods FastSpeciesTree (VeryFastTree & IQ-TREE), Phyling (FastTree & IQ-TREE), busco2phylo-nf and FFP compared to a set of reference *Opsithokont* phylogenies.

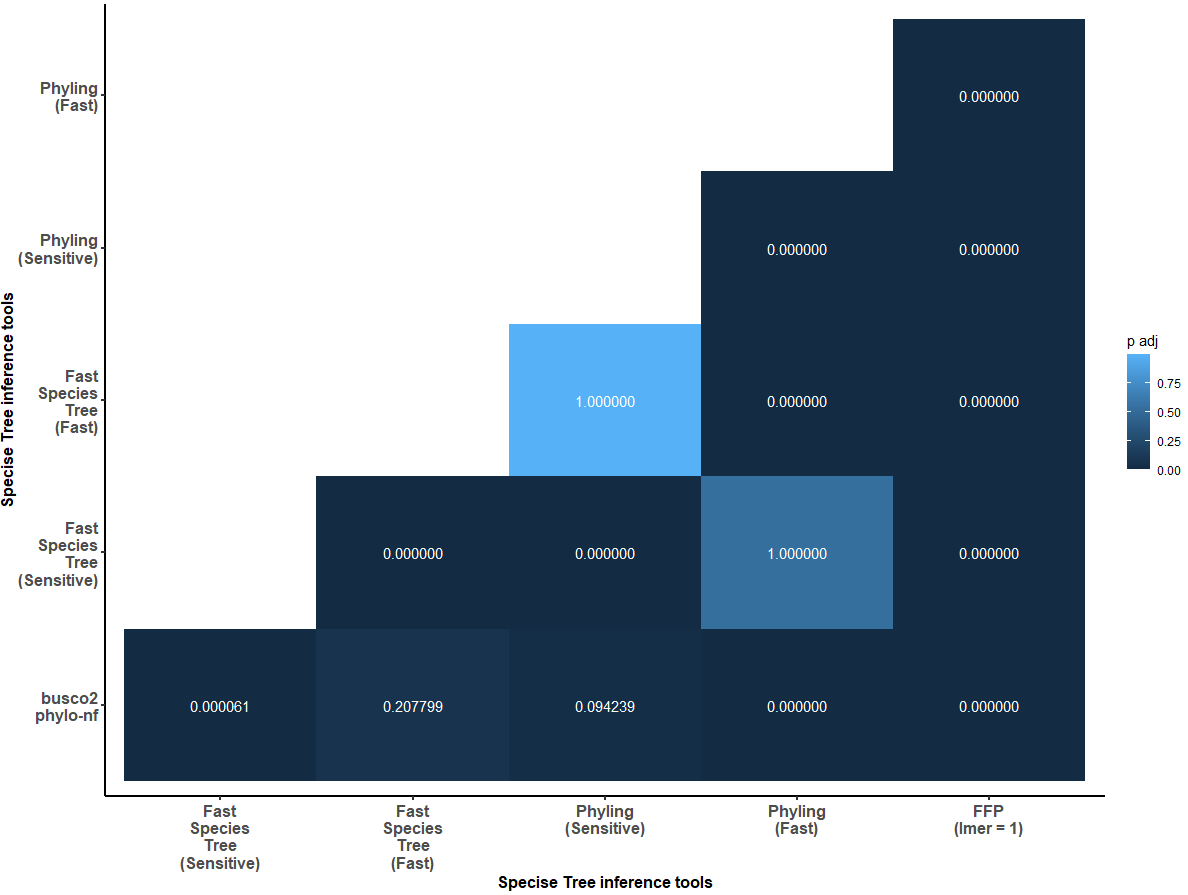

**Figure 2.** The statistical similarity of Quartet distance between 4 species tree inference methods FastSpeciesTree (VeryFastTree & IQ-TREE), Phyling (FastTree & IQ-TREE), busco2phylo-nf and FFP compared to a set of reference *Opsithokont* phylogeny.

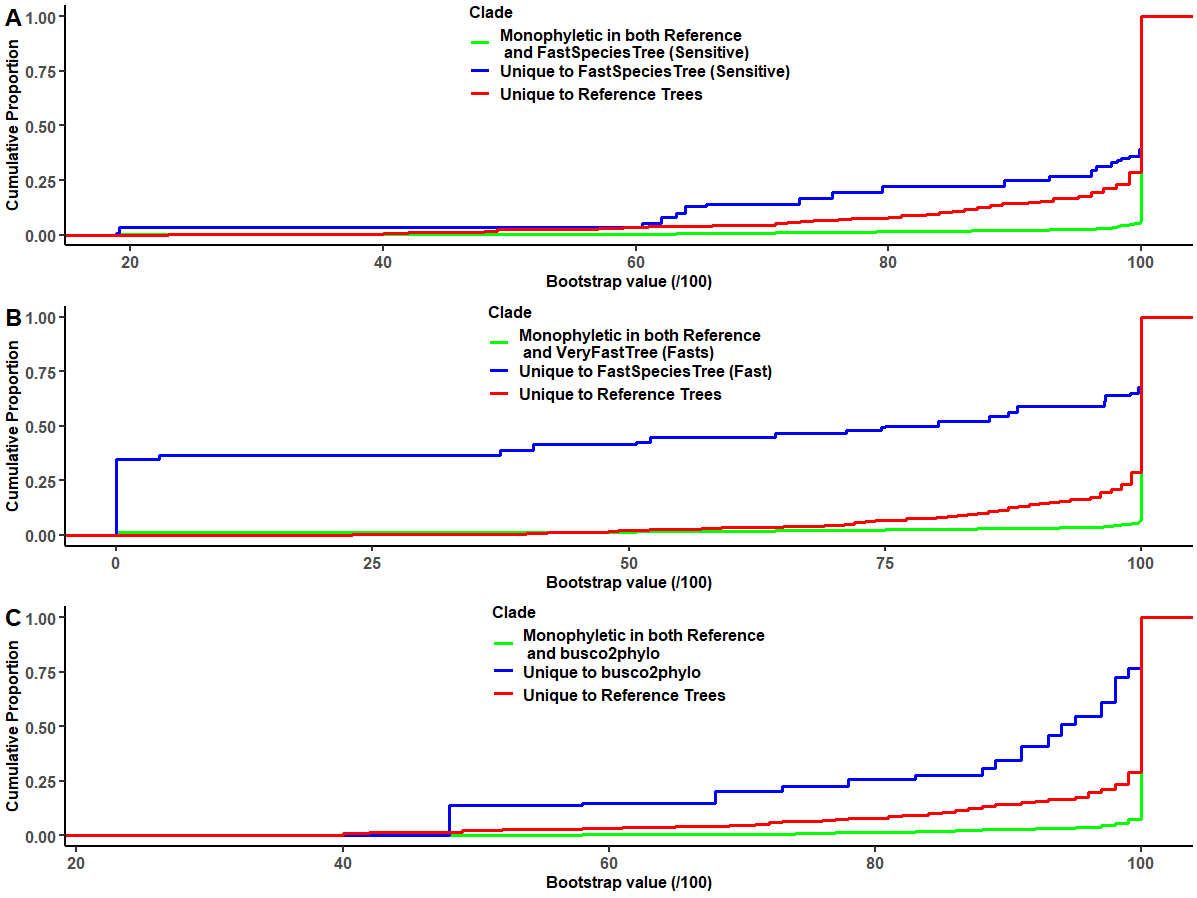

**Figure 3.** Support (bootstrap) values for monophyletic clades shared (red) or unique to the reference (green) or method (blue) for FastSpeciesTree with IQ-Tree (A), VeryFastTree (B) and busco2phylo-nf (C).

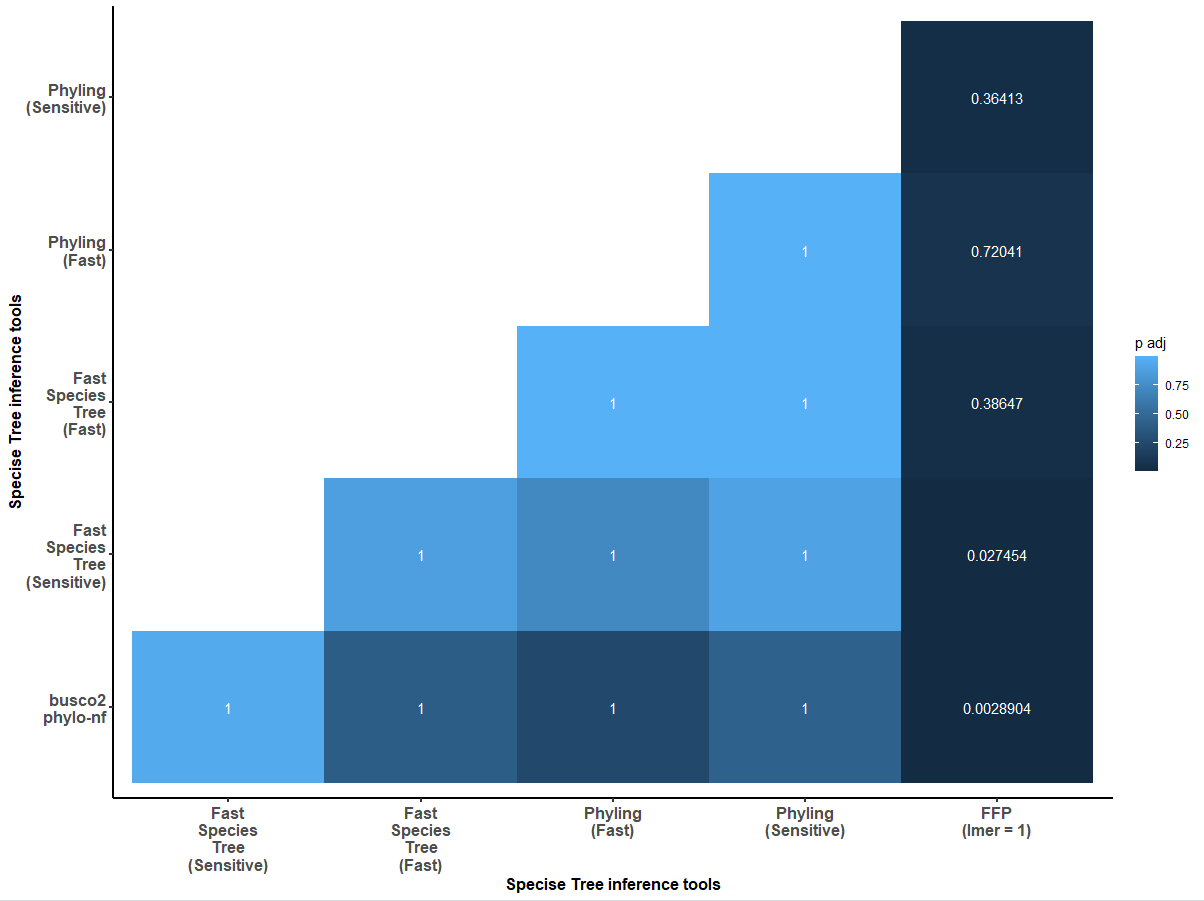

**Figure 4.** The statistical similarity of generalised Robinson-Foulds distance e between 4 species tree inference methods FastSpeciesTree (VeryFastTree & IQ-TREE), Phyling (FastTree & IQ-TREE), busco2phylo-nf and FFP compared to a reference phylogeny of budding yeast.

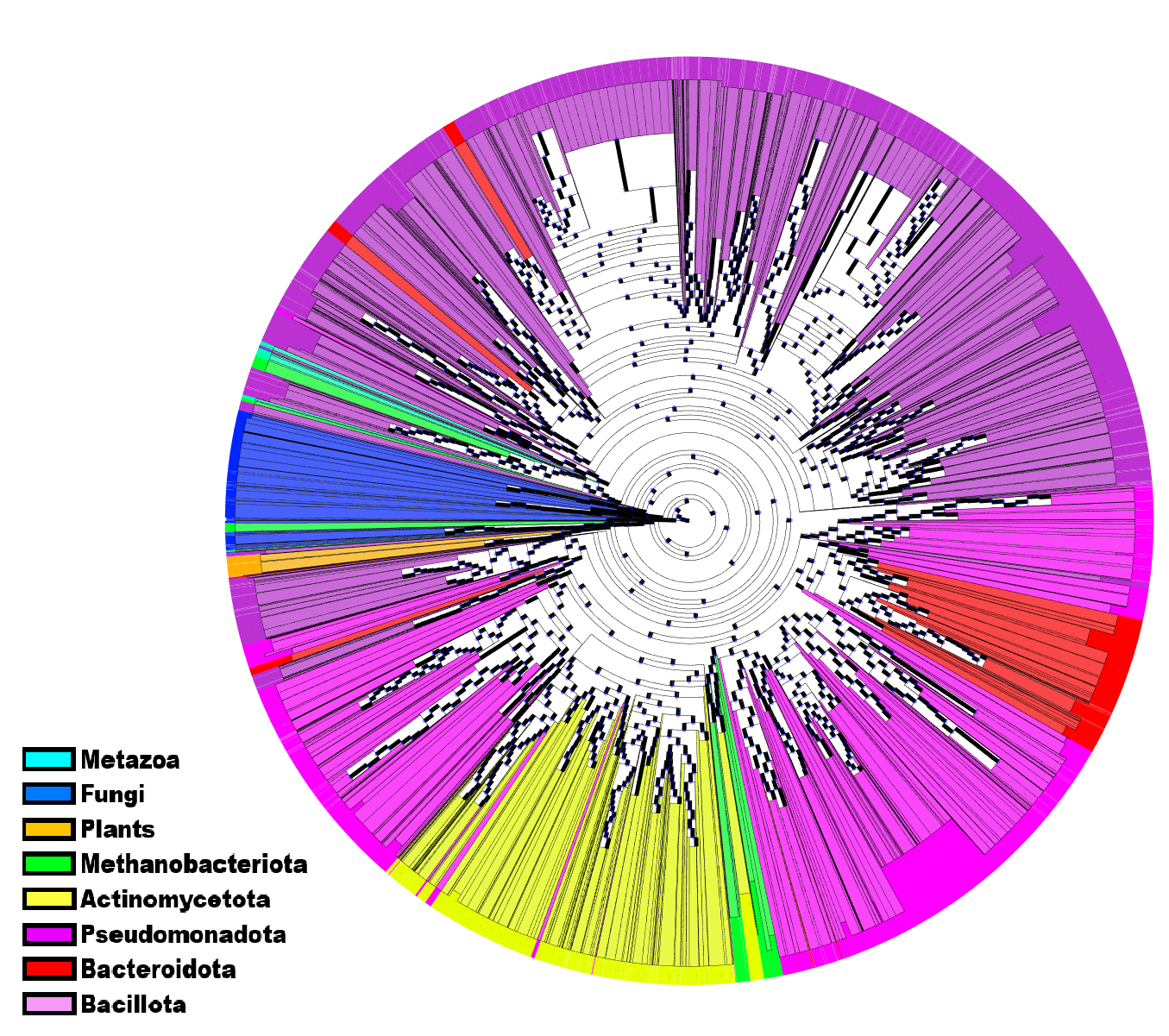

**Figure 5.** A phylogeny of 28,244 species containing Eukaryotes and Prokaryotes from ensemble generated using Phyling with 20 concatinated genes from the Eukaryote BUSCO schema. Major bacterial phyla along with Eukaryotes are highlighted. Annotations for the plants, fungi and metazoa are based on the ensembl dataset. Bacteria are clustered into the major phyla of *Actinomycetota, Psuedomonadota, Bacillota, Bacteroidota* and *Methanobacyeriota* using BacDive. The figure is visualised using ete4.

**Table 1.** The proportion of taxa from a particular group (major phyla or domain of life) that can be found on a single clade which contains at least 95% of that group from the ensemble tree of life generated by FastSpeciesTree and Phyling from 28,244 proteomes.

|  | FastSpeciesTree | Phyling |
| --- | --- | --- |
| Metazoa | 100 | 31.69398907 |
| Pseudomonadota | 99.98526161 | 85.40664545 |
| Fungi | 99.85915493 | 97.59206799 |
| Plants | 98.61111111 | 98.49246231 |
| Methanobacteriota | 100 | 91.14688129 |
| Actinomycetota | 99.89255976 | 97.24919094 |
| Bacteroidota | 99.65397924 | 67.0471464 |
| Bacillota | 99.9526216 | 35.70405728 |
